# PennZzz - an algorithm for estimating behavioral states from wrist-worn accelerometery

**DOI:** 10.1101/347807

**Authors:** Richard McCloskey, Matthew H. Goodman, Sara McHugh Grant, Arpita Ghorai, Philip R. Gehrman, Maja Bućan

## Abstract

Sleep is a heterogeneous behavioral state comprised of different stages and interspersed with episodes of wakefulness. Sleep/wake states can be monitored in the sleep laboratory by polysomnography (PSG). However, sleep studies are intrusive, laborious and expensive, and are usually performed over a single night. In contrast, wrist-worn activity-tracking devices (actimeters) are inexpensive, unobtrusive, and can be used to estimate sleep and wake patterns over multiple nights. We designed the PennZzz algorithm to estimate sleep and wake from actimetry data. Results obtained by actimetry-based monitoring in 26 subjects were compared to stages of sleep and wakefulness detected by simultaneous polysomnography. We found that our algorithm identifies PSG-defined wake episodes with a high accuracy (336/431 – 76% of algorithm wake events correspond to true wakefulness). Furthermore, we find that the algorithm is sensitive enough to detect the majority (258/431 – 59%) of true wake episodes occurring after the first NREM1 to NREM2 transition. With correction, algorithm outputs can be used to estimate the total amount of time awake after sleep onset. We further refined this program for application in a high-throughput manner to assess the total amount of sleep, wake, and non-wear during longer recording periods.

## Introduction

Sleep is a heterogeneous process comprised of multiple behavioral states including REM, NREM, and wake episodes (Zepelin et al., 2005). Overnight polysomnography (PSG), the “gold standard” assessment of sleep, requires monitoring of electroencephalogram (EEG), electrooculogram (EOG) and electromyogram (EMG). The use of in-lab PSG for sleep assessment is limited due to cost, reactivity to sleeping in a novel laboratory environment, and ability to record over a single or small number of nights. In contrast, activity data obtained using wrist-worn accelerometer monitoring (i.e. actimetry) has been used for the past few decades as a proxy measure of sleep (Sadeh, 2011; Meltzer et al., 2012). Actimetry devices are inexpensive, can be worn unobtrusively on the body, and in addition to an accelerometer, can contain other biosensors such as light, temperature and heart rate monitors. As shown for several research-grade devices, the patterns of activity can be used to infer the subject’s sleep/wake state using validated algorithms that have demonstrated good agreement with PSG (Sadeh et al., 1994). These devices can be worn for several weeks to months, enabling long-term monitoring of activity and sleep patterns in a naturalistic environment.

Activity monitors, also referred to as actigraphs, have traditionally measured movement in a single axis; however, a new type of device utilizes microelectromechanical systems (MEMS) that record changes in acceleration in three axes, and as such, are more sensitive to a wider variety of movements and velocities (te Lindert et al 2013). For example, the recently introduced GENEActiv Sleep device (Activinsights, Kimbolton, UK) contains sensors to record acceleration, temperature, and light (Zhang et al., 2012; Pavey et al., 2014; Scott et al., 2017) and is less expensive than traditional research grade devices. While MEMS devices provide more in-depth activity assessment, there is a paucity of validated algorithms for the analysis of patterns of physical activity or sleep/wake states.

We sought to expand accelerometry data analysis by developing an analytical program flexible enough to accurately assess patterns of both activity and sleep. Using GENEActiv devices, we measured activity patterns and developed PennZzz - an algorithm that can detect: a) sleep within active phase; b) activity during sleep phase (waking events); and c) behavioral states such as light, intermediate and deep sleep. To validate these traits, we correlated detected behavioral states with PSG-assessed sleep stages. We show that our algorithm can be used to assess a number of sleep parameters including the amount of wake after sleep onset, a trait known to be associated with poor sleep quality. Lastly, we applied the algorithm in a high-throughput way to sequentially analyze sleep patterns over multiple days to facilitate downstream analysis of overall sleep trends in a large number of subjects.

## Material and Methods

### Subjects

The study consisted of two parts, a two-week naturalistic actigraphy portion and a single night polysomnography (PSG) assessment. For the naturalistic portion of the study, two types of subjects were included. 100 participants (ages 10-40) were recruited through the Pennsylvania Longitudinal Study of Parents and Children (PALSPAC) twin registry and through advertising in the Philadelphia metropolitan region. Additionally, patients with a history of affective disorders were recruited from an existing patient registry at the University of Pennsylvania Mood and Anxiety Disorders Program. 26 subjects were included in the PSG study. All procedures were approved by the University of Pennsylvania Institutional Review Board and all subjects provided written informed consent prior to participation in the study.

### Actigraphy

All subjects were instructed to wear the GENEActiv device for two consecutive weeks in the naturalistic portion of the study. Subjects in the PSG portion were instructed to wear the device the day before, day of, and day after their sleep center visit. The GENEActiv Sleep device recorded acceleration (g-force), temperature (°C), and light (lux) in 30 datapoints per 1 second (30 Hz). Data were then converted to 1-second epochs using the GENEActiv 2.9 Software (Activinsights, Kimbolton, UK).

### Polysomnography

Standard polysomnographic procedures were used to record the EEG, EOG, EMG, and EKG using the Sandman System. Electrode placements of FpZ, CZ, OZ were used according to the International 10/20 system. Two EOG electrodes were placed, positioned 1 cm below and lateral to the outer canthus of the left eye and 1 cm above and lateral to the outer canthus of the right eye. Two surface EMG electrodes were taped onto the chin 2 cm apart. Additional leads were used to measure leg movements and breathing. Two electrodes were taped over the anterior tibialis muscle of each leg to detect leg movements during the night. Flexible Resp-EZ belts were placed around the abdomen and chest to measure breathing-related movements during the night. A nasal cannula was used to detect pressure and an oximeter probe was placed on the finger to measure blood oxygen saturation.

### Data Analysis

Acceleration data were transformed into the sum vector magnitude:

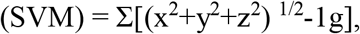

which corresponds to the sum of acceleration in three dimensions with gravity subtracted, and is further comprised of the sum of all readings in the designated 1 second epochs. The <u>t</u>empo<u>v</u>ascillitory <u>a</u>ctivity (TVA) was then calculated by finding the variance in SVM in a 121-second moving temporal window surrounding the 1-second epoch of interest. If fewer than 121 datapoints were present because the moving window was at the beginning or end of a dataset, then that window was truncated down to a minimum of 61 data points. The program then thresholded TVA into one of five separate states: WAKE, DEEP SLEEP, INTERMEDIATE SLEEP, LIGHT SLEEP and ambiguous. The ambiguous TVA values, which were greater than the threshold for identifying light sleep but less than the threshold for identifying wake, were identified as WAKE or SLEEP if the epochs immediately surrounding the ambiguous epoch were identified as WAKE or SLEEP, respectively. Additionally, absolute temperature was used to assess whether individuals may have removed their watch. In instances when behavior resembled sleep, a temperature check was also performed, and when individuals displayed very low movement TVA matching deep sleep, and when the absolute temperature was below 24°C, behavior was classified as NON-WEAR (Fig. 6). Note that this method of assessing NON-WEAR was not validated, except for a few tests on field data, and that very cold individuals who are asleep may register with a NON-WEAR score, even when they are wearing the device.

We also calculated variance in temperature within a 181-second temporal window (the <u>t</u>empo<u>v</u>ascillitory <u>t</u>emperature (TVT)), which was calculated in a similar manner as TVA except 181s window was used, and a single threshold bisects a bi-modal distribution observed in a log transformation of the TVT data. This threshold determines whether the device is in or out of a homothermic state. TVT is not directly used in the program to assess behavior, but is considered a unique trait on its own that might inform results in future studies since TVT is higher during sleep than during wakefulness.

To compare algorithm outputs to corresponding sleep stages, PSG data were generated in 30s epochs, and were then aligned to match algorithm outputs based on the time recorded by the computer. Data were then compared pairwise at each 1s epoch and summed to calculate the correlation between the state determined from actimetry data and the state determined by PSG.

We assessed the relationship between algorithm WAKE scores and true PSG-measured wakefulness in the following 3 ways. 1) we calculated the sensitivity of the algorithm to detect true wake episodes (detection of a true wake episode by both algorithm and PSG), 3) we measured the accuracy of the algorithm (Percentage of total WAKE episodes by algorithm that correspond to a PSG-defined wake episode), and 3) compared the total WAKE amount scored by algorithm to the wake amount scored by PSG. For the sensitivity and accuracy measurements we restricted analysis to the time period after the individual first transitioned out of NREM1 up until wakefulness. This time frame was chosen because computational detection of short wake episodes was error-prone during the beginning NREM1 phase of sleep since limb movements are higher during this stage (data not shown). Therefore, our analysis ignores any WAKE episode detection during the portion of sleep prior to the first transition out of NREM1.

To execute these three analyses we, with computer assistance, aligned and co-plotted algorithm and PSG data, then applied a set of rules to score correlations as follows to score sensitivity and accuracy of wake episodes in sleep as follows: We defined a true wake episode as any single un-interrupted consecutive period of time where the independently scored PSG registered wakefulness. Then, for each instance in which the algorithm registered WAKE, but no PSG data registered wake, the result was scored as a false positive episode. If PSG data registered wake with no WAKE episode registering by algorithm, then the score was false negative. If both PSG and algorithm registered wake at the same instance, then the score was a true wake episode. If multiple algorithm WAKE episodes were scored during one single PSG-defined wake episode, those multiple algorithm WAKE episodes were each counted as “true positive” to calculate accuracy, but were counted as a single true positive to calculate sensitivity. The reciprocal situation of detecting one single algorithm WAKE episode while multiple PSG-defined wake episodes occurred was more rare, but was scored in a similar manner. A separate measurement method was used to compare the absolute quantification of total time in WAKE scored by actimetry *vs.* wake by PSG. For this comparison, the total number of minutes scored as WAKE by the algorithm was directly compared to the total minutes of wake scored by PSG.

To perform the weekend trait analysis, data from individuals wearing the GENEActiv for between 10-14 days without more than 4 days of non-wear were selected. Each day was then processed with the PennZzz algorithm and data were exported into .csv formats and imported into Microsoft Excel. Dates were then manually entered by referencing the raw data output from GENEActiv, and the weekday or weekend mean of traits was assessed.

### Software development

To train and program an algorithm for GENEActiv data that might detect sleep stages, we initially generated images displaying sleep stages as defined by polysomnography of one subject. Then, we visualized raw outputs for GENEActiv on the same time scale with similarly sized images to examine whether GENEActiv outputs could be correlated to sleep stages. On visual inspection of the figures, we observed that GENEActiv SVM measurements drift when limb movement was low, as for example happens during sleep. In contrast, TVA values were far more stable during immobility and therefore better predictors of the sleep state. We therefore examined trends using TVA. We examined 2-3 arbitrary TVA thresholds to see if they could differentiate WAKE, LIGHT SLEEP, and DEEP SLEEP stages. Threshold values were deemed useful if they divided behavioral states that correlated well with those states determined polysomnographically. We performed further algorithm training on the data obtained from three additional subjects.

TVA thresholds varied slightly for different individuals, perhaps owing to device or individual differences. To control for this inter-individual variability and automate the identification of TVA thresholds appropriate to a particular individual, we wrote a program to find the peak in the histogram distribution of log transformed TVA data, and designates this as the LIGHT SLEEP threshold. The software then uses this threshold to select the INTERMEDIATE SLEEP threshold positioned automatically between the DEEP and LIGHT thresholds. The DEEP and WAKE thresholds are fixed values. The original algorithm development was performed using MATLAB (Mathworks Inc), and was later translated to Python, and expanded to be able to analyze more than 20 hours of data per algorithm cycle, and to automatically step through GENEActiv .csv output files for each day in the dataset.

## Results

All subjects were instructed to wear the GENEActiv device for two consecutive weeks. A separate set of subjects in the PSG portion of the study were instructed to wear the GENEActiv device the day before, day of, and day after their sleep center visit. The GENEActiv Sleep device recorded acceleration (g-force), temperature (°C), and light (lux) in 30 datapoints per 1 second (30 Hz). These recorded data were downloaded from devices using GENEActiv software (Activinsights Ltd., Kimbolton, UK) and exported into a comma separated value (csv) file in 1 second epochs for analysis by the PennZzz software. Figure 1 displays the three data outputs from GENEActiv device.

**Figure 1.**
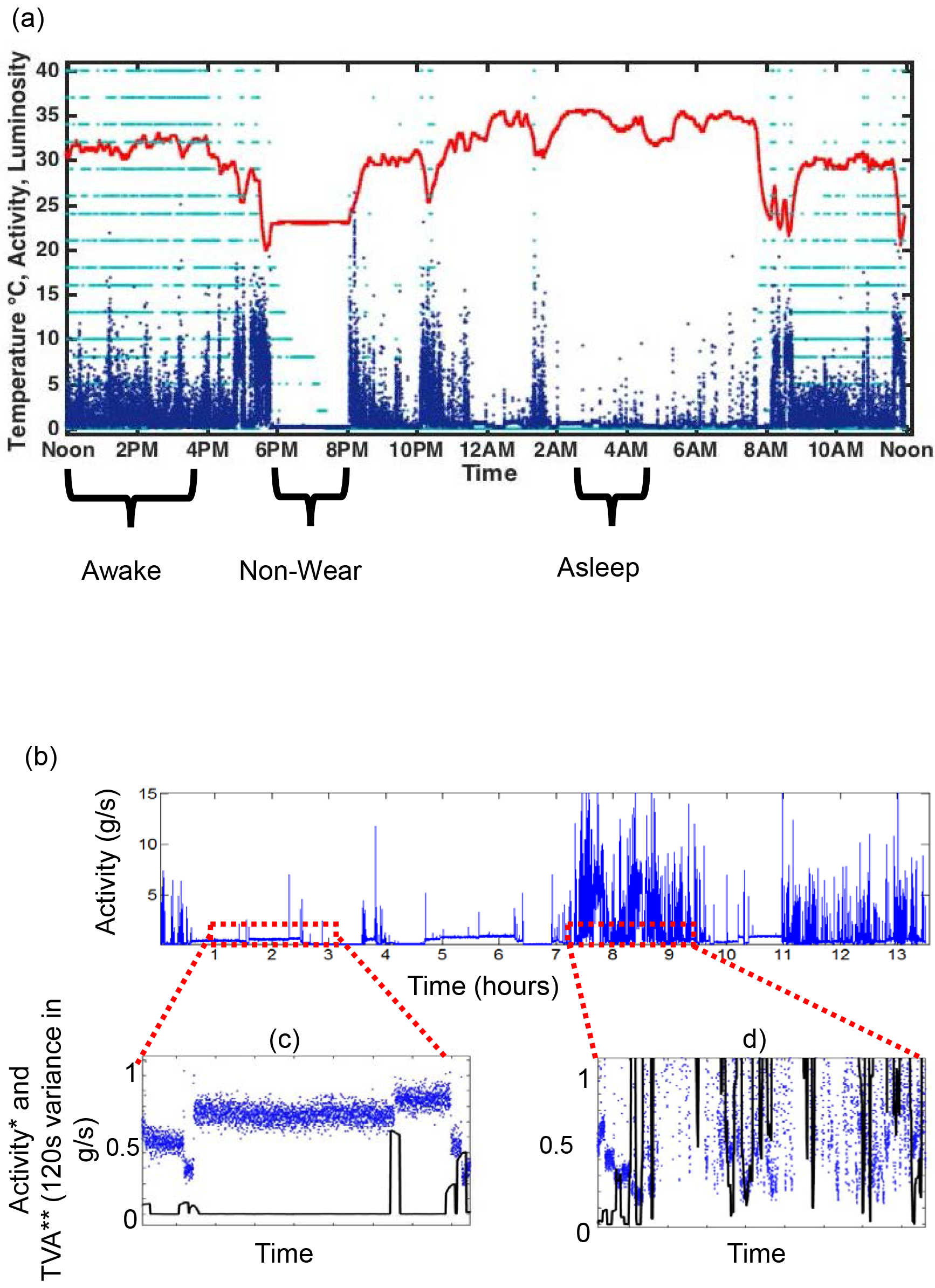
Accelerometer outputs illustrating variance as a method to monitor sleep and wake. (a) Raw data from a GENEActiv 1.2 device worn for 24 hours; Red = Temperature (°C); Navy Blue = acceleration in Gs; Teal = Lux; (b) The raw activity values measured by GENEActiv devices (blue line plot); Values are generally lower when individuals are sleeping (c, blue dots) when compared to wake (d, blue dots), but baseline drift in values is evident. In contrast, tempovascillitory activity (TVA) yields low values during sleep (b, black line) regardless of activity drift, and yields generally higher values during wake (c).

**Table 1.** Example comparison of select traits comparing weekdays or weekends. To assess whether our algorithm could infer meaningful data about human behavior in an ethological setting we analyzed data from 10 individuals with our algorithm. By briefly glancing at graphical outputs of each day manually we discarded days containing nonwear in excess of 4 hours, high levels of non-wear undetected by the algorithm, and days in which the beginning or end of the rest period appeared to be wrong by greater than ½ an hour. From this data we were able to analyze 127 days, and we further divided days in to weekdays (Sun-Thurs) or weekends (Fri-Sat). We selected interesting traits that appeared to differ as expected or that did not. Our analysis confirms past results that individuals sleep more on weekends, primarily attributed to late awakening times. Attributes such as sleep efficiency do not differ between weekdays and weekends, but some attributes, such as the amount of time spent homothermic, do.

| **Trait** | **Weekday\* Value** | **Weekend Value** |
| --- | --- | --- |
| Subjects (n) | 10.00 | 10.00 |
| Days analyzed (n) | 90.00 | 37.00 |
| Total days of sleep (h) | 7.62 | 8.62 |
| Light and intermediate sleep (h) | 6.60 ± 0.20 | 7.70 ± 0.34 |
| Deep sleep (h) | 1.00 ± 0.10 | 1.00 ± 0.10 |
| Bedtime | 11:17 PM | 11:35 PM |
| Wake time | 7:02 AM | 8:43 AM |
| Sleep in rest period only (h) | 5.70 ± 0.20 | 6.70 ± 0.30 |
| Algorithm WASO (m) | 95.8 ± 3.80 | 113.2 ± 17.40 |
| Predicted real WASO (m) | 64 | 76 |
| Algorithm sleep efficiency | 78 ± 1% | 77 ± 1% |
| Predicted real sleep efficiency | 84% | 84% |
| Mean daily activities (g) | 1.42 ± 0.03 | 1.24 ± 0.05 |
| Mean activity during wake (g) | 1.80 ± 0.03 | 1.63 ± 0.07 |
| Mean activity during rest period (g) | 0.61 ± 0.02 | 0.61 ± 0.03 |
| Homothermic state during sleep (h) | 0.07 ± 0.03 | 0.27 ± 0.08 |

The sum vector magnitude (SVM) values, calculated based on each of the three dimensional accelerometer axes (see Material and Methods), are generally lower during sleep than during WAKE, yet at some points indistinguishable from SVM values during WAKE (Fig. 1), at times when the subject was awake yet not moving. We therefore examined whether the variance of SVM values in a 121-second moving window, the tempovascillitory activity (TVA; Fig. 1) may better differentiate sleep and behavioral inactivity during wakefulness.

We observed that the log distribution of TVA values for SLEEP-WAKE was more useful for distinguishing behavioral states as data largely distributes into a relatively clear distribution with at least two modes (bimodal) (Fig 1). When confirmed by quantitative comparison we observed log-transformed TVA values were significantly higher during wakefulness than during sleep states, indicating a higher level of variability in movement during waking (Fig. 1). Moreover, deeper sleep states such as NREM2 and NREM showed overall lower log(TVA) values than light NREM1 sleep stage (NREM1 differed from REM, NREM2, and NREM3, *p<0.02* in each case) (Fig. 2b). log (TVA) values were not different between REM sleep and NREM2 or NREM3 even though REM sleep is characterized by higher brain arousal, perhaps because muscle atonia of REM sleep prevents limb movement.

**Figure 2.**
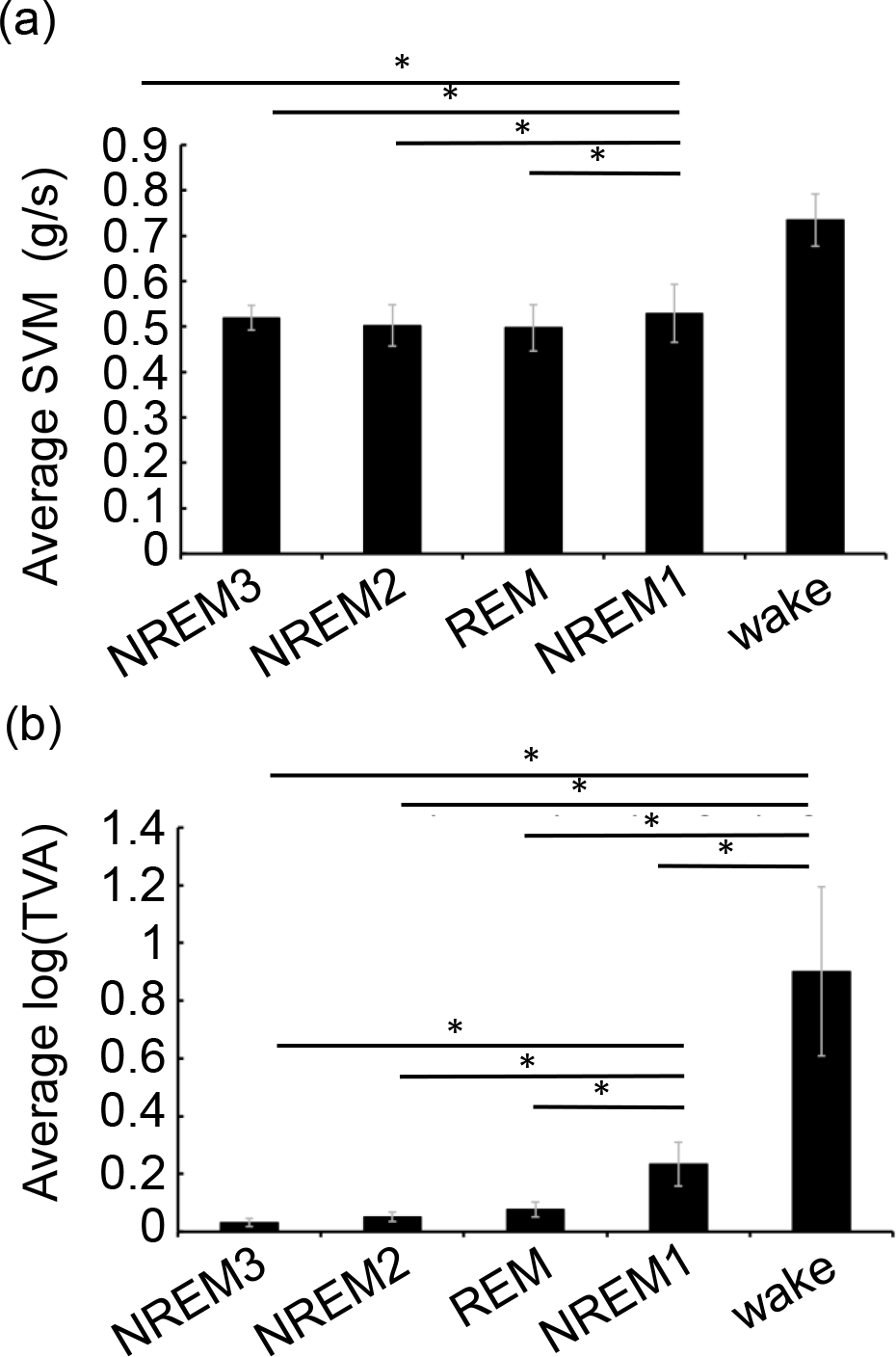
Wake and NREM1 sleep stages are associated with higher variation in activity values than other sleep stages. 26 subjects spent a night in the sleep lab after which PSG-derived sleep stages were compared to GENEActiv data. (a) Average activity per second based on the corresponding sleep stage. (b) Tempovascillitory activity (TVA) calculated by finding the variance in 121-second moving windows. Error bars denote standard error of the mean (SEM) of 26 individual subjects. (a, b) Asterisks mark p<0.02.

An alternative approach to compare PSG-defined sleep stages with actimetry-based traits such as total activity and TVA is to first distinguish between SLEEP and WAKE and then define the composition of each state in terms of PSG stages. To take this actimetry-based approach, TVA was stratified into separate bins corresponding to a) DEEP SLEEP; b) INTERMEDIATE SLEEP; c) LIGHT SLEEP and d) WAKE. To determine how these states could be delineated we observed the distribution of TVA values during sleep (Fig. 3a). We manually chose two thresholds (Fig. 3b-c), which determined the points above and below which WAKE and SLEEP were probable, respectively. These original thresholds were chosen by visually comparing the SLEEP/WAKE status of a single subject (unpublished data). We then designed the program to choose the threshold below which sleep is considered INTERMEDIATE, and a second threshold, below which sleep is considered DEEP, by finding the local maximum in log(TVA) data corresponding to sleep. Simple visual comparison of PennZzz outputs with PSG recorded sleep stages suggested that PennZzz can accurately identify waking epochs and, in addition, can stratify sleep into DEEP SLEEP and LIGHT SLEEP. (Fig. 3d).

**Figure 3.**
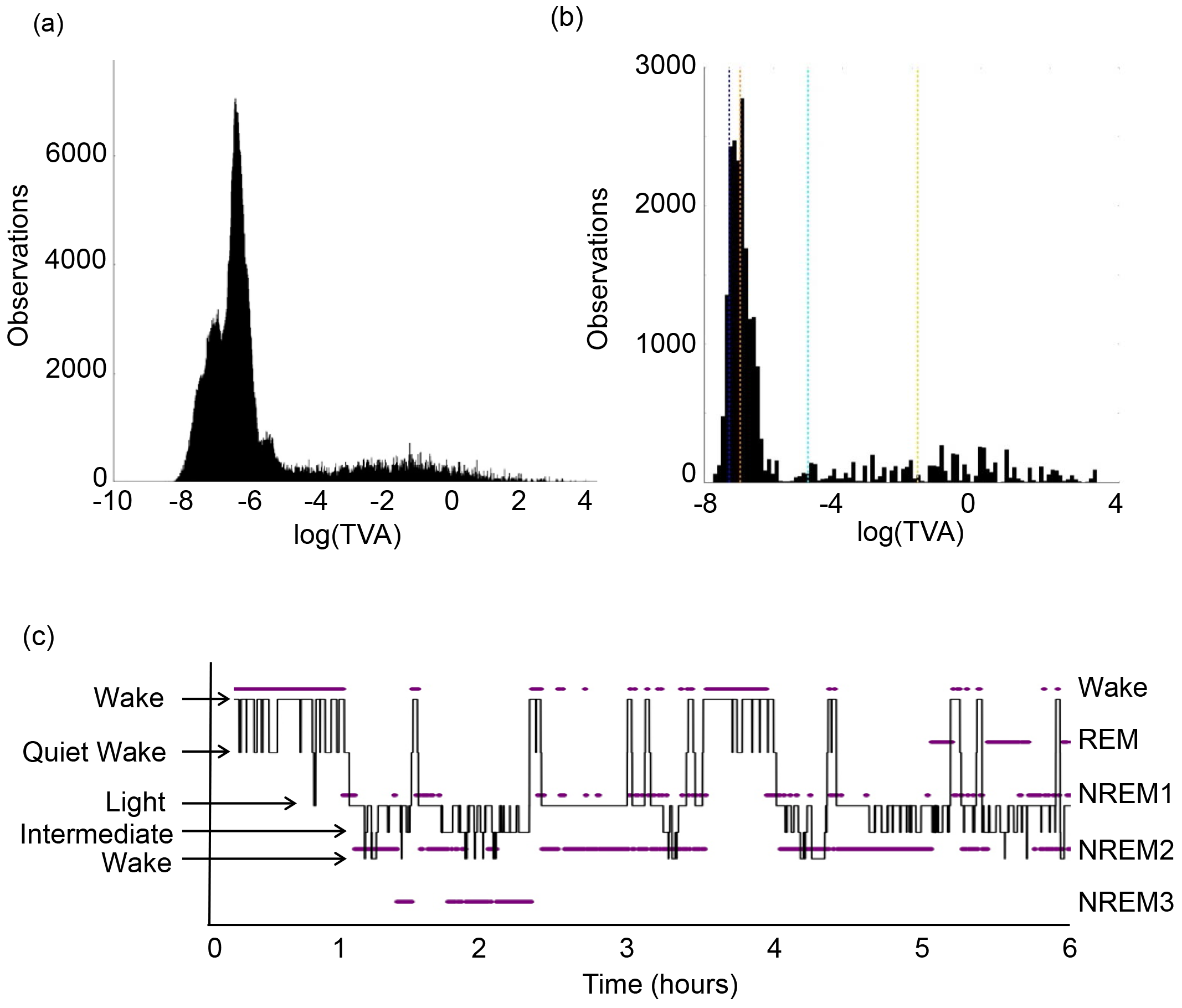
A method to parse algorithm behavioral states from TVA values. (a) The distribution of log transformed TVA data corresponding to the night rest period of 26 subjects monitored in the sleep lab. Low values (left) represent lower amounts of GENEActiv TVA corresponding to less movement while high values (right) correspond to higher TVA. (b) Graphical representation of arbitrary behavioral states chosen by algorithm parameters to divide sleep behavior into <u>A</u>lgorithm determined behavioral states. Behavior is segmented into bins based on the vertical dotted lines shown on this plot. Starting from the left-most vertical line, data to the left of each line can be described as follows: Blue- flagged as deep sleep, Orange- flagged as intermediate sleep, Teal- flagged as light sleep, Yellow- flagged as unknown and assigned by the algorithm based on neighboring behavior, and lastly, to the right of the Yellow line behavior is called wake. (c) Graph describing how TVA thresholds are chosen, and which <u>A</u>lgorithm determined <u>b</u>ehavioral <u>s</u>tate scores are assigned. A temporal alignment of sleep stages (purple dot scatter) with <u>A</u>lgorithm determined <u>b</u>ehavioral <u>s</u>tate scores (continuous black line plot).

We compared PennZzz-defined behavioral states to PSG-defined sleep stages. Figure 4 shows the probability of classifying each sleep stage correctly using PennZzz. PSG-defined sleep stages NREM2 and NREM3 were more closely correlated to DEEP and INTERMEDIATE SLEEP scores than to LIGHT SLEEP or WAKE as defined by PennZzz. Conversely, PSG-defined wake and NREM1 were more likely to be correlated to WAKE and LIGHT SLEEP states as defined by PennZzz (*p < 0.005* when comparing the theoretical frequency of state distribution correlations to the actual distribution). PSG-defined REM sleep epochs were most likely to be classified as light sleep but with less specificity than NREM1 or wake. This observation suggests that deeper sleep “stages” are correlated to lower Algorithm determined behavioral state scores and therefore lower TVA values. However, in a second comparison, we excluded PSG wake data from the theoretical distribution model to test whether sleep stages still stratify across ABS states in a statistically significant way, but we found no significant correlation (*p*=0.45). This second test shows that while there is a trend for Algorithm behavioral state stages to correspond to certain sleep stages, the difference is not robust. Overall, these data demonstrate that actimetry-based TVA thresholds can define behavioral states that probabilistically correspond to specific PGS-detected sleep stages.

**Figure 4.**
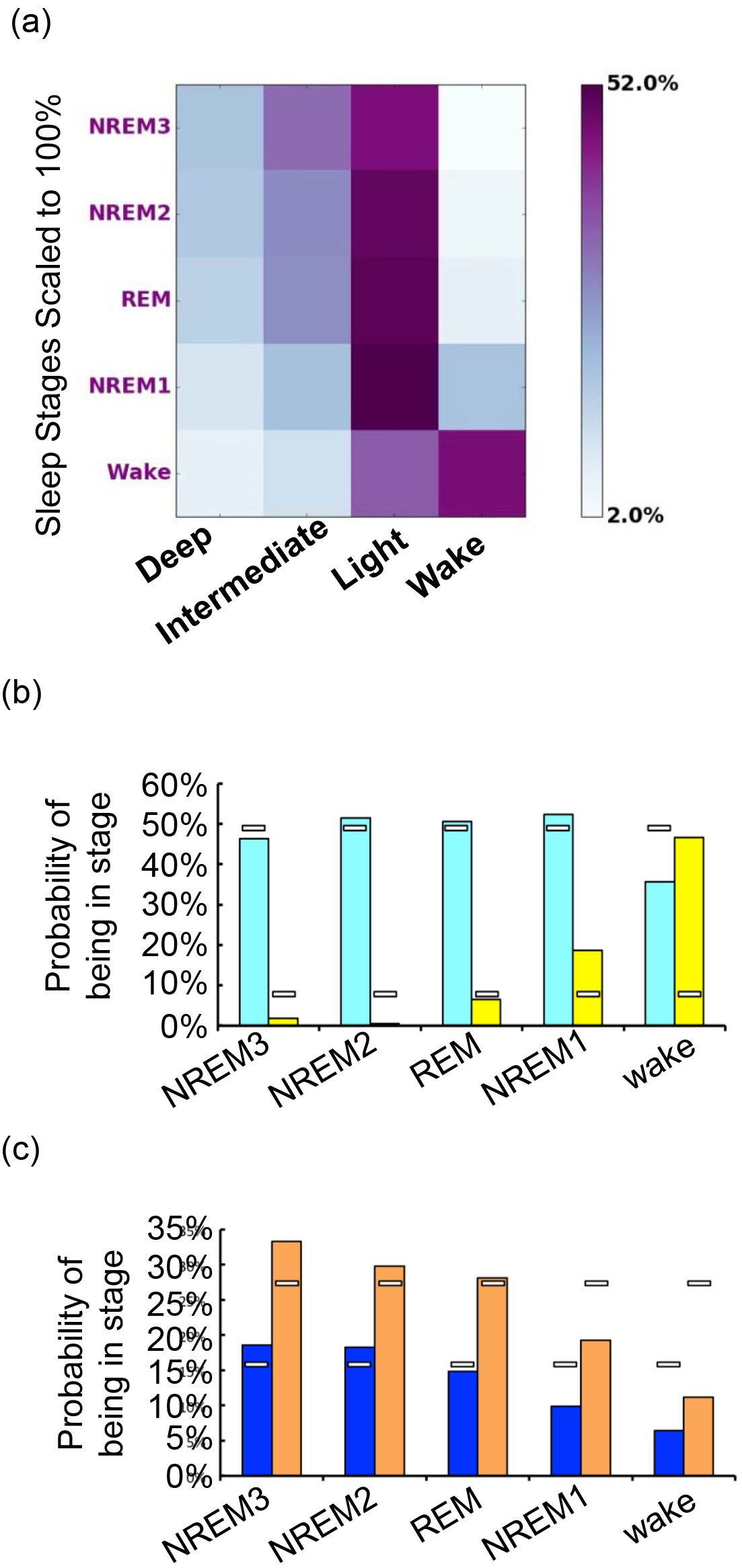
Algorithm behavioral state scores probabilistically correspond to sleep stages. (a) A heat map showing the distribution of sleep stages across <u>A</u>lgorithm determined <u>b</u>ehavioral <u>s</u>tates. Each sleep stage (rows) in the colormap was normalized to 100% for 26 individuals. The distribution of sleep stages stratifies across Algorithm behavioral states in a way significantly different than expected showing <u>A</u>lgorithm determined <u>b</u>ehavioral <u>s</u>tates probabilistically correlate to the likelihood of a sleep stage. chi^2^ 12df, N=26)= 63.5 *p<0.005.* Most of the algorithm predictive power is due to strong wake detection since, without wake, the distribution shift trend is insignificant *(p=0.45)*.a (b) Probability of being assigned the indicated <u>A</u>lgorithm determined <u>b</u>ehavioral <u>s</u>tates score (light and wake) for each sleep stage (x axis). White bars mark the expected height if the null hypothesis is true. (c) Similar depiction for deep and intermediate Algorithm behavioral state scores. For each sleep stage in (b and c) bars sum to 100%.

High levels of wake after sleep onset (WASO) are believed to be disruptive to sleep; therefore, the ability to quantify WASO accurately could be useful in assessment of sleep quality. We examined whether PennZzz can estimate the timing and amount of WASO. We aligned the sleep stages assessed by PSG to Algorithm behavioral state scores for each subject (Fig. 5a), and then assessed how often PSG and Algorithm behavioral state scores agreed on whether subjects were in a wake episode (see Material and Methods) (Fig. 5b) by manually counting episodes that occurred after the first onset of sleep. We found that, after accounting for false positives and false negatives, the algorithm was able to correctly identify 59% of all wake episodes scored by PSG. Without counting false negatives, 76% of all episodes defined as Algorithm behavioral state WAKE correlated to a true wake episode on PSG showing relatively high accuracy (Fig. 5b). Overall, these results show that the program correctly identifies the majority of true waking events during sleep. We next analyzed actimetry patterns to quantify the total amount of time that the algorithm scored as WAKE, beginning at the time of lights-off recorded on the PSG, and compared this to PSG time scored as awake. There was a linear relationship between the PSG vs. Algorithm behavioral state scored wakefulness with an R^2^ val.=0.5 (Fig. 5c). In general, these data show that the algorithm overestimates the total time spent awake per episode (Fig. 5c-d), but that values can be corrected by linear regression analysis (equation displayed in Fig. 5c) to be closer to a true value. Lastly, we compared the rate of concordance at each instant as to whether the actimetry-based algorithm and PSG generally agreed on the status of being awake or asleep, and found that for the first 5 hours after lights-out, the agreement rate is 95.4% (Fig. 5e).

**Figure 5.**
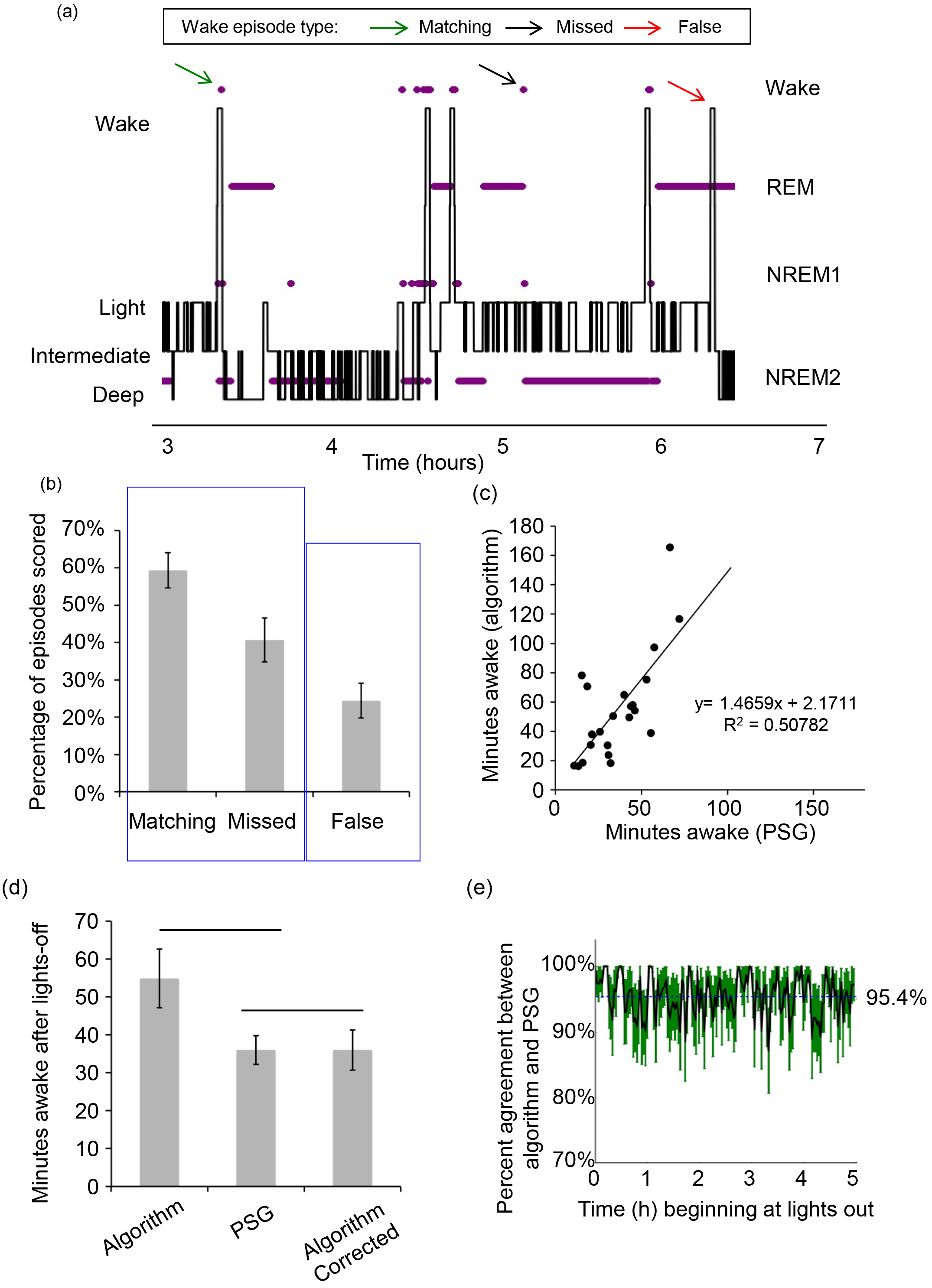
Wake episodes and total wake after sleep onset can be estimated by algorithmically detecting waking events. (a) A temporal alignment of four sleep stages (purple dot scatter) with Algorithm behavioral state scores (continuous black line plot). A green arrow, black arrow, and red arrow point out examples of a positively identified wake episode (matching), a missed wake episode, and a falsely identified wake episode respectively. (b) Quantification of three categories of wake episodes shows that a majority of true wake episodes can be detected by Algorithm behavioral state scoring. (c) Regression analysis of Algorithm behavioral state determined WASO amount compared to PSG assessed WASO. The data suggests that <u>A</u>lgorithm determined <u>b</u>ehavioral <u>s</u>tates scoring can be used to calculate WASO, and that the Algorithm behavioral state scoring model accounts for about 50% of the variance (R^2^=0.5) in predicting WASO. *n=21* because individuals with r >80 minutes of wake as assessed by PSG were excluded. (d) Algorithm behavioral state scoring for wake significantly overestimates *p=0.003* the total WASO, but can be corrected by assuming the linear model. (e) Algorithm and PSG agreement in 10 minute bins during the first 6 hours of the rest period as beginning at the time of “lights off’ in the sleep lab. A black line plots % agreement and green bars show shows SEM for *n=26* individuals.

The PennZzz analyses described above compared activity-based and PSG-detected traits only at night. In order to facilitate the PennZzz analysis of activity and sleep patterns over the 24-hour day (or several days), we used the algorithm to identify the main sleep episode. Specifically, we identified the first instant at which the next 3 hour span of time was scored as 70% sleep and the time point at which 70% of the next 2 hour span corresponds to wakefulness. Thus, the PennZzz analysis of actimetry traits can robustly identify typical consolidated rest periods for most of the subjects in this sample (Fig. 6).

**Figure 6.**
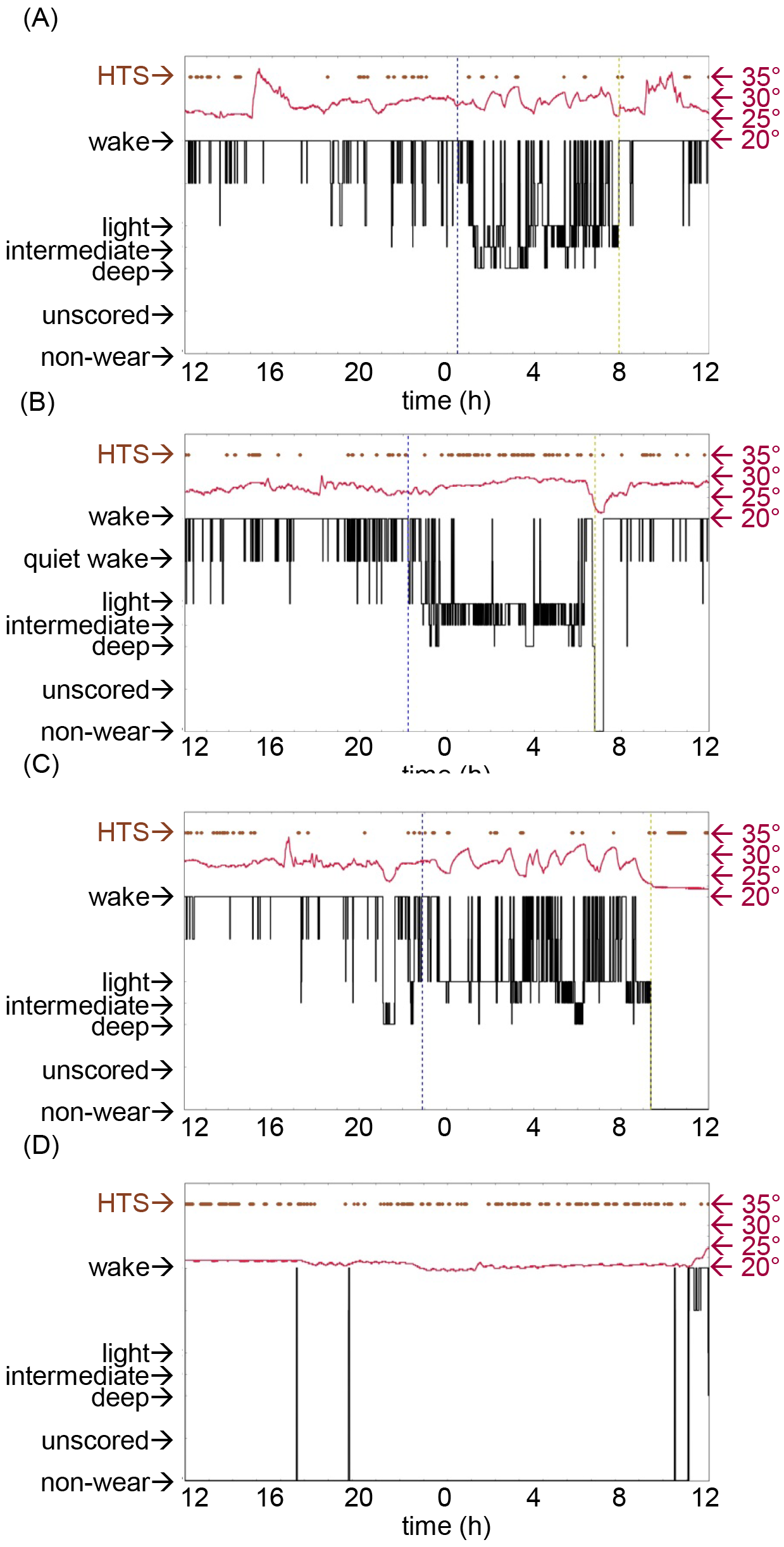
Detection of homothermic states or low absolute temperatures is an efficient pre-condition to detecting GENEActiv device non-wear. (a-d) Each plot shows a data over the course of a 24 hour period of wear for one subject beginning at 12 noon, and ending at 11:59:59 AM the following calendar date. Multiple data types are shown: **Algorithm behavioral state** score (black line), absolute temperature (crimson line), temperature-stable homothermic states (brown dot scatter), the algorithmically assessed beginning of the main rest period (blue dashed vertical line), and the algorithmically assessed end of the main rest period (yellow dashed vertical line). (a) A representative individual’s behavior on the day and night prior to their sleep study (b) the same individual’s behavior on the day of their sleep study. (c) The individual’s behavior the following day, which includes detection of an extended period non-wear initiated after waking. (d) GENEActiv device was unworn for the whole day.

Finally, in order to differentiate DEEP SLEEP, marked by low TVA, from “NON-WEAR” we combined the actimetry data with temperature recordings collected by the GENEActiv, reasoning that a more uniform environment, i.e. cooler and more stable temperatures would indicate that devices are not worn. We observed changes in absolute temperature and also calculated variance (or oscillations) in 181s window time periods throughout the day to identify homothermic states. Initially, we scored as NON-WEAR all windows when temperature fell below 24°C (Fig. 6). Moreover, a comparison between actimetry and removals recorded on the sleep diary showed that these states are marked by a less variable homothermic phase, i.e. low tempovascillitory temperature (TVT) (Fig. 7a). In order to further confirm this in field data, we also examined temperature and TVT values for individuals for whom we had PSG data by examining the temperature in the controlled environment of the sleep lab. We found that NREM3, NREM2, and REM registered as having both higher absolute temperatures and more stable temperature compared to wakefulness (Fig. 7b-c). Thus, our algorithm is reliable in identifying instances of NON-WEAR by a combination of detecting temperature stability and absolute temperature. In some cases this method is sensitive enough to detect NON-WEAR just before or after the main rest episode. Somewhat unexpectedly, we observed that temperature homothermic states and the absolute temperature measured by devices are also elevated during sleep.

**Figure 7.**
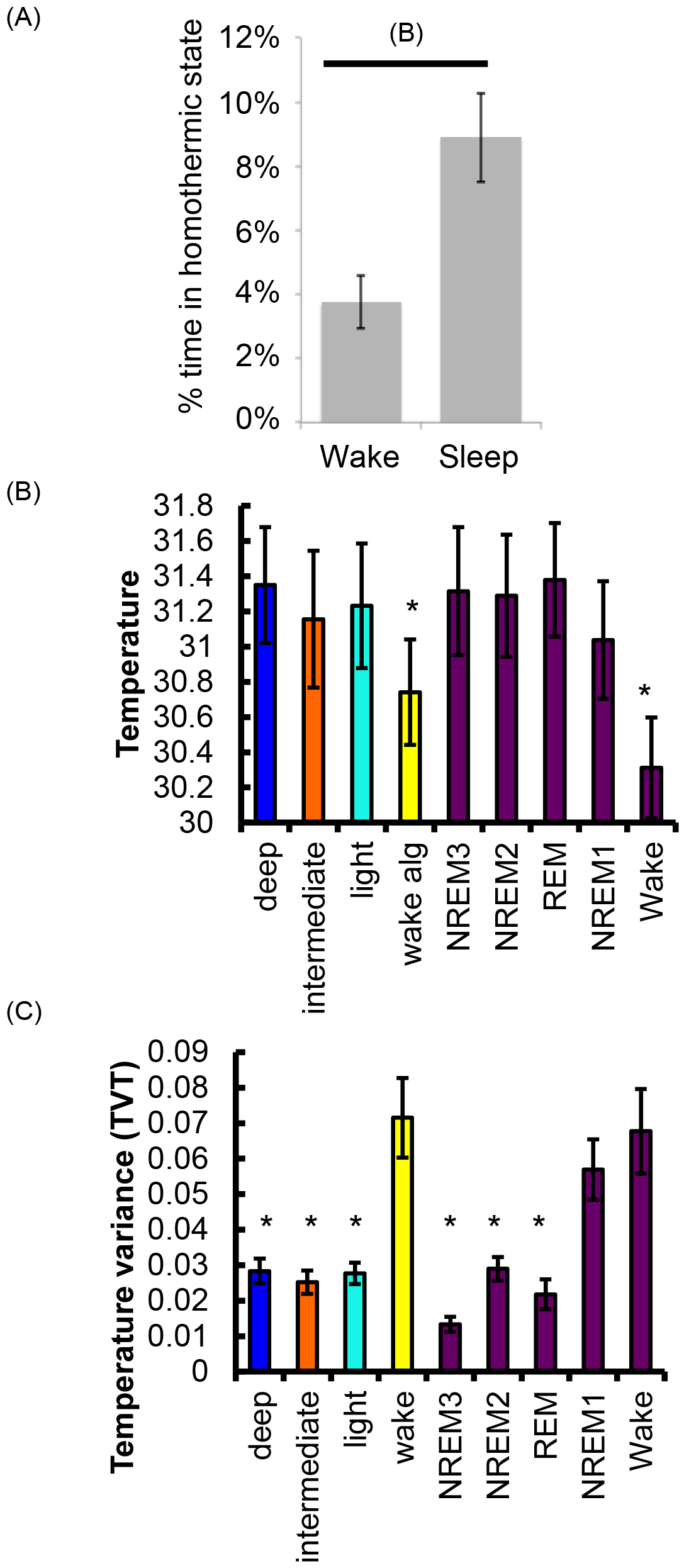
GENEActiv temperatures are warmer and more stable during sleep. (a) The data for 45 days across 6 subjects was analyzed, and the total amount of time spent in a homothermic state when in the main rest period or not was calculated. The data show that during the main rest period a larger proportion of time is spent in a homothermic state indicating temperatures are less variable. (b) We examined data for the 26 recorded nights in the sleep lab and observed absolute temperatures differ based on the sleep stage and ABS stage. In particular, absolute temperatures are lower when individuals are awake. (c) Tempovascillitory temperature (TVT) was calculated for each ABS score and sleep stage.

## Discussion

Our study describes the development and validation of an algorithm for the analysis of sleep and wake traits based on tri-axial actimetry data collected using GENEActiv Sleep, a research-grade wrist-worn accelerometer. By combining patterns of movement with temperature, the PennZzz algorithm can detect with reasonable accuracy: a) sleep within the active phase; b) activity during the sleep phase (waking events); and c) behavioral states such as LIGHT, INTERMEDIATE and DEEP SLEEP.

The PennZzz algorithm takes a unique approach to estimating SLEEP and WAKE by computing second-by-second variability in movement, in contrast to prior algorithms that have largely used a weighted moving window that estimates SLEEP and WAKE by examining the activity level in a given epoch as well as several of the preceding and following epochs (Cole et al., 1992). While other methods have been proposed, most rely on the absolute activity level in each epoch. This is the first algorithm, to our knowledge, that uses variation in activity over time to aid in distinguishing behavioral states.

Our PennZzz algorithm can distinguish different behavioral states within sleep. We call the algorithm states we used to describe sleep either LIGHT, DEEP, or INTERMEDIATE, which are each characterized by different patterns of variation in activity. It is tempting to identify these labels with traditional PSG-defined sleep stages. Indeed, we find that REM sleep, correlates to our LIGHT SLEEP state. However, the comparison with PSG shows that there is not a high correspondence between these behavioral states and sleep stages. Future algorithm refinement may lead to greater ability to differentiate NREM from REM sleep using actimetry data.

In recent years, there has been an explosion in commercially available actimetry devices that are marketed to consumers, mainly to monitor activity, but also to measure sleep quality and sleep amount (Wright et al., 2017). There is tremendous interest in the value that these devices might have for conducting research studies. Like the present algorithm, the proprietary software outputs of these devices stratify sleep into arbitrary assigned states based on the measurement of movement, often giving these states names like “light, deep, restful,” *etc.* However, device manufacturers rarely share details of the algorithms used to identify behavioral states, nor do they make raw data available, making it difficult to evaluate the validity of manufacturer claims. For the time being, researchers will need to rely on devices such as the GENEActiv.

This study provides a description and initial validation of the PennZzz algorithm, but there is much more work to be done. There is a need to validate the algorithm in independent samples with a range of sleep patterns. The sample in this study consisted of generally healthy good sleepers. A weakness of prior algorithms is that they have lower agreement with PSG in individuals with insomnia (Lichstein et al., 2006). This is likely due to the fact that individuals with insomnia lie in bed still but awake and this inactivity is scored as sleep. Future studies are needed to examine the utility of this algorithm in poor sleepers to see whether it suffers from the same limitations or if the higher resolution, tri-axial data are more sensitive to fine movements that would identify these periods of quiet wakefulness. This study was also limited in sample size, so the collection of additional data will help with future validation. Finally, this study only collected PSG data during the night so it was not possible to validate estimates of sleep and wake during daytime hours. A successful algorithm needs to be able to accurately differentiate wake and sleep at all times of day and night.

Going forward, our tool will be useful for investigators that have collected GENEActiv data in assessing WASO, daily activity patterns, and temperature characteristics during sleep and wake. With further refinement, the PennZzz algorithm will likely be applicable to data collected with other MEMS-type actimeters. Actimetry has proven to be useful for both research and clinical purposes, and a new generation of devices and scoring algorithms will likely expand the accuracy and range of uses in the future.

## Acknowledgments

We thank David Raizen, Chris Fang-Yen and Nalaka Goonerante for discussions in the early stages of the project and comments on the manuscript. We thank Holly Barilla, Emma Greger and Allison Jacoby for help with the recruitment. R.M., P.R.G. and M.B. designed the study, R.M. and A.G. developed the algorithm, M.H.G. and S.M.H. recruited study subjects, R.M., A.G. performed data analysis and R.M., P.R.G. and M.B. wrote the manuscript.

